# Range contractions of the Broad-winged Hawk in the Northeast United States

**DOI:** 10.1101/2021.10.06.463411

**Authors:** Rachael Pruitt, Laurie Goodrich, Matthew B. Shumar, Andrew M. Wilson

## Abstract

The Broad-winged Hawk (BWHA, *Buteo platypterus)* is a small, secretive hawk with distinguishing broad black tail bands that breeds in northeastern North America. The hawk nests in deciduous or mixed forest, often near water, and close to clearings or forest edges. Land conversion and fragmentation alters the landscape and reduces the area of contiguous forest used by BWHA. This study seeks to determine the landscape characteristics influencing the apparent breeding range declines of the BWHA at the landscape scale. Landscape characteristics and BWHA presence data from 18,684 Breeding Bird Atlas blocks (each about 25km^2^) from Ohio, West Virginia, Maryland, Pennsylvania, and New York for two atlas period (1st Atlas: 1980s, 2nd Atlas: 2000s) were analyzed. Bayesian latent Gaussian models were fitted using INLA to determine best fit model for predicting the landscape characteristics associated with BWHA presence. The best models included landscape changes in land cover, including forest, water, urban, barren, farmland, and wetland and fragmentation of the landscape. Trends in loss were especially prevalent around the region’s largest cities: New York, Philadelphia, Baltimore and Washington DC. Loss of BWHA at the block-level was associated with areas with less forest in the 2000s, a decline in size of largest forest patches, lower elevations and lower latitudes. We suggest that both habitat loss and climate change may be contributing to the range contraction of the Broad-winged Hawk in the northeast United States.

## INTRODUCTION

Habitat loss and fragmentation are threats to biodiversity throughout the world and are of utmost concern to wildlife conservationists (Fahrig 2003, 2013; Haddad et al. 2015). Habitat fragmentation is defined as the division of habitat into several isolated patches, resulting in a reduction of suitable habitat for any given species (Rolstad 1991). The formation of edges, especially in forest habitats, increases inter-specific competition, changes the structure of available habitat, opening up niches for some species but leading to a loss in the area of suitable habitat for forest-specialist species, particularly those that avoid forest edges.

Forest cover in the Northeastern United States has partially recovered from wholesale deforestation following European colonization. Large patches of contiguous secondary forest in the Appalachian and Adirondack Mountains support a high diversity of bird species, making them an important region for examining the impacts of land use change on bird distributions (Abrahams et al. 2015). However, in areas peripheral to the mountains, lower elevation forests are highly fragmented, with conversion to urban development a prevalent cause of forest loss and fragmentation (Drummond and Loveland 2010). The Northeast United States contains some of the largest cities in the country, including the Northeast Megalopolis, which contains almost one-fifth of the country’s population (Kulkarni 2015). As these urban centers grew, the urban and exurban expansion resulted in extensive urban sprawl, which continues to replace forested and natural lands. Estimates of forest loss in the region between 1973 and 2000 total 10.05 million ha, with a net decline in forest cover of more than 3.70 million ha (Drummond and Loveland 2010). In addition to exurban sprawl, a recent additional driver of forest fragmentation in the Northeast United States, particularly in Pennsylvania, is the development of infrastructure for oil and natural gas extraction. Exploration for development sites is becoming more prevalent throughout the Allegheny Plateau of Pennsylvania, as well as nearby West Virginia and Ohio (Drohan et al. 2012). In Pennsylvania, 45% of existing pads are located within forests, of which 23% are in core forest (Drohan et al. 2012), with rates of loss of privately-owned core forest more than twice of public forests (Langlois et al. 2017).

The breeding range of the Broad-winged Hawk (*Buteo platypterus*) extends from Nova Scotia south through the Appalachian Mountains and peripheral low-elevation forests into northern Florida and west into eastern Oklahoma (Goodrich et al. 2014a). The northern extent of the range extends west through the Great Lakes region and northwest through southern Alberta (Goodrich et al. 2014b). Broad-winged Hawks most often nest in deciduous or mixed forest, often near water, and close to small clearings or forest edges (Fitch 1974, Titus and Mosher 1981, Crocoll and Parker 1989, Goodrich et al. 2014a). The close proximity to water is most likely due to the higher prey density found near water sources (Titus and Mosher 1981, Crocoll and Parker 1989). Additionally, forest edges and upland openings are thought to be used as primary hunting sites for the Broad-winged Hawk due to prey availability and accessibility (Rosenfield 1984).Thus, some degree of habitat heterogeneity may be beneficial to this species. However, forest fragmentation can lead to a decrease in availability of small vertebrate prey, the main food sources of the Broad-winged Hawk (Fitch 1974, McCabe et al. 2019). At the local scale, these characteristics are important for nest site selection, but less is known about how habitat extent and configuration at the landscape scale influence Broad-winged Hawk nesting densities, productivity, and population trends. It is assumed that contiguous forests provide the best landscape for nest sites of the Broad-winged Hawk (Goodrich 2012), but to what extent this species is able to persist in highly fragmented forests is not known. Determining whether land cover change has an impact on the breeding range of this species will be valuable information for better understanding its ecology and conservation needs.

Statewide breeding bird atlases have indicated localized losses of Broad-winged Hawks in Maryland (Ellison 2010), New York (McGowen and Corwin 2008), Pennsylvania (Goodrich 2012), Ohio (Rodewald et al. 2016) and West Virginia (Bailey and Rucker 2021). Many of these decreases are thought to be associated with increasing fragmentation and urbanization (Goodrich 2012). Unlike some species of hawks, the Broad-winged Hawk is rarely found in small woodlots or in built-up landscapes, therefore competition with larger hawk species for nesting sites may increase as suitable habitat becomes less available (Ellison 2010), and this small *buteo* may be more affected by the higher nest predation rates associated with more fragmented forests (Thompson III et al. 2002). While Broad-winged Hawks are currently listed as least concern by the IUCN, they continue to face pressure from climate change and land conversion (BirdLife International 2012). Preemptive understanding of how habitat change in the Northeast United States is impacting the Broad-winged Hawk could be beneficial for understanding and managing the habitats of this and other forest hawk species. This study uses bird atlas data to examine the landscape characteristics that are associated with local changes in the breeding range of the Broad-winged Hawk in the Central Appalachian Mountains, Adirondack Mountains and surrounding areas of the Northeastern United States. We hypothesize that forest fragmentation is the main driver of the observed loss of Broad-winged Hawks from some areas since the 1980s.

## METHODS

### Breeding Bird Atlas Data

Our study uses block-level Broad-winged Hawk presence data from the breeding bird atlases of five contiguous states in the Northeastern United States: Maryland (including the District of Columbia), New York, Ohio, Pennsylvania, and West Virginia. Breeding bird atlases are useful for monitoring range shifts of birds because they are repeated approximately every 20 years, and provide data at large spatial scaled (Dickinson et al. 2010). The atlases used in this study are based on approximately 5×5 kilometer (c. 3×3 mile) gridded sampling blocks. The study area encompasses 18,684 atlas blocks, equaling 471,590 km^2^. Sampling for each atlas was first conducted during the 1980s and repeated during the 2000s (Table 1). Hereafter, we refer to these time periods as 1st Atlas and 2nd Atlas respectively. Breeding bird atlas data were collected primarily through volunteer effort, using standardized procedures. The protocols ideally ensure consistency in coverage among blocks and requires that a significant amount of time is invested in each sampled block (Porter and Jarzyna 2013). Although atlas methodologies are designed to ensure consistent coverage, in reality, coverage is inconsistent, both spatially (between blocks) and temporally (between atlas projects) (Wilson et al. 2017).While measures of effort can be used to account for such variation, it is important to note that effort information, such as number of visits, search effort (e.g. total hours), and years of visits are not available for all ten atlas projects used in our analysis.

**Table 1.** State bird atlas projects data used for analysis including date ranges and number of blocks included.

| State | No.<br>blocks | 1st Atlas years (citation) | 2nd Atlas years (citation) |
| --- | --- | --- | --- |
| Maryland/DC | 1,294 | 1983-87 (Robbins et al. 1996) | 2002-06 (Ellison 2010) |
| New York | 5,332 | 1980-85 (Andrle et al. 1988) | 2000-05 (McGowen and Corwin 2008) |
| Ohio | 4,437 | 1982-87 (Peterjohn et al. 1991) | 2006-11 (Rodewald et al. 2016). |
| Pennsylvania | 4,937 | 1985-89 (Brauning et al. 1992) | 2004-09 (Wilson et al. 2012) |
| West Virginia | 2,653 | 1984-89 (Buckelew and Hall 1994) | 2009-14 (Bailey and Rucker 2021) |

### Landscape and Land Cover Metrics

We used ArcMap 10.3.1 (ESRI, Redlands, CA) to calculate landscape metrics within each atlas block for both atlas periods. We used land cover data (National Land Cover Database) for the period closest to the midpoint of the atlas sampling period for the 2nd Atlas (MD, NY, PA: 2006; WV, OH: 2011) (Fry et al. 2011, Homer et al. 2015) and 1992 land cover data for the 1st Atlas, (the earliest land cover data available at these scales). The National Land Cover database uses a 16 land cover classification scheme at a spatial resolution of 30 meters for 2001 and later. Land cover data for 1992 uses a 21 class system at a 30 meter spatial resolution. Because 1992 land cover data uses a 21 class system at a 30 meter spatial resolution, which were not compatible with more recent land cover products, we used the retrofit land cover change product to determine 1992 land cover values to maintain a comparable classification scheme (Fry et al. 2009). We tabulated the area of each of seven broad categories of land cover within each block: water, urban, barren land, forest, shrubland/grassland, farmland, and wetlands; and calculated the Shannon diversity index of the seven broad land cover types within each block (Flather and Sauer 1996).

To obtain block-level metrics of forest fragmentation we used the Landscape Fragmentation Tool (LFT) geoprocessing package (Shapiro et al. 2016), which uses morphological image processing to differentiate forest land cover into core and edge classifications (Vogt et al. 2007).The LFT identifies four forest classes: 1) core forest that does not have non-forest boundaries, 2) patch forest which is forest that is too small to contain core forest, 3) perforated forest which comprises the boundaries between core forest, and 4) small patches of non-forest and edge forest which are the boundaries of forest surrounding large areas of non-forest (Vogt et al. 2007). The four categories determined by the LFT were then grouped into two forest classes: 1) core and perforated, and 2) patch and edge. Tests of the tool revealed that 7×7 cell neighborhoods resulted in forest edge bands of approximately 90 meters, most consistent with a frequently used definition of core forest being at least 100 meters from a forest edge. The area for each of the three defined classes (non-forest, core, and edge) was tabulated for each block. The number of core forest patches per block, and the area of the largest patch of core forest that was intersected by the block were calculated. The core to edge ratio, a commonly employed measure of forest fragmentation, was determined by dividing the area of core by the total area of forest (Imre 2006).

We derived topographical data for each block from a digital elevation model (DEM; USGS, http://nationalmap.gov/elevation.html) and calculated mean, maximum, minimum and range of elevation. The latitude of each block centroid was also calculated in ArcGIS.

### Statistical Analysis

Because survey coverage was variable, both spatially and temporally, and detection of birds is a function of observer effort, we included a proxy of survey effort as a covariate in our models (Sadoti et al. 2013). Actual survey effort (e.g. total hours spent in each block) was not available for every atlas dataset, so we used recorded species richness as a proxy for effort. Our proxy measure for survey effort is a ratio between number of species observed in the atlas block and a uniform standard of 75 species to represent block completion (henceforth “sp.ratio”). In doing so, we assume that the probability of detecting Broad-winged Hawks in an atlas block is linearly correlated with the number of species detected up to 75 species, but that this relationship plateaus for blocks with more than 75 species detected (Figure 1).

**Figure 1.**
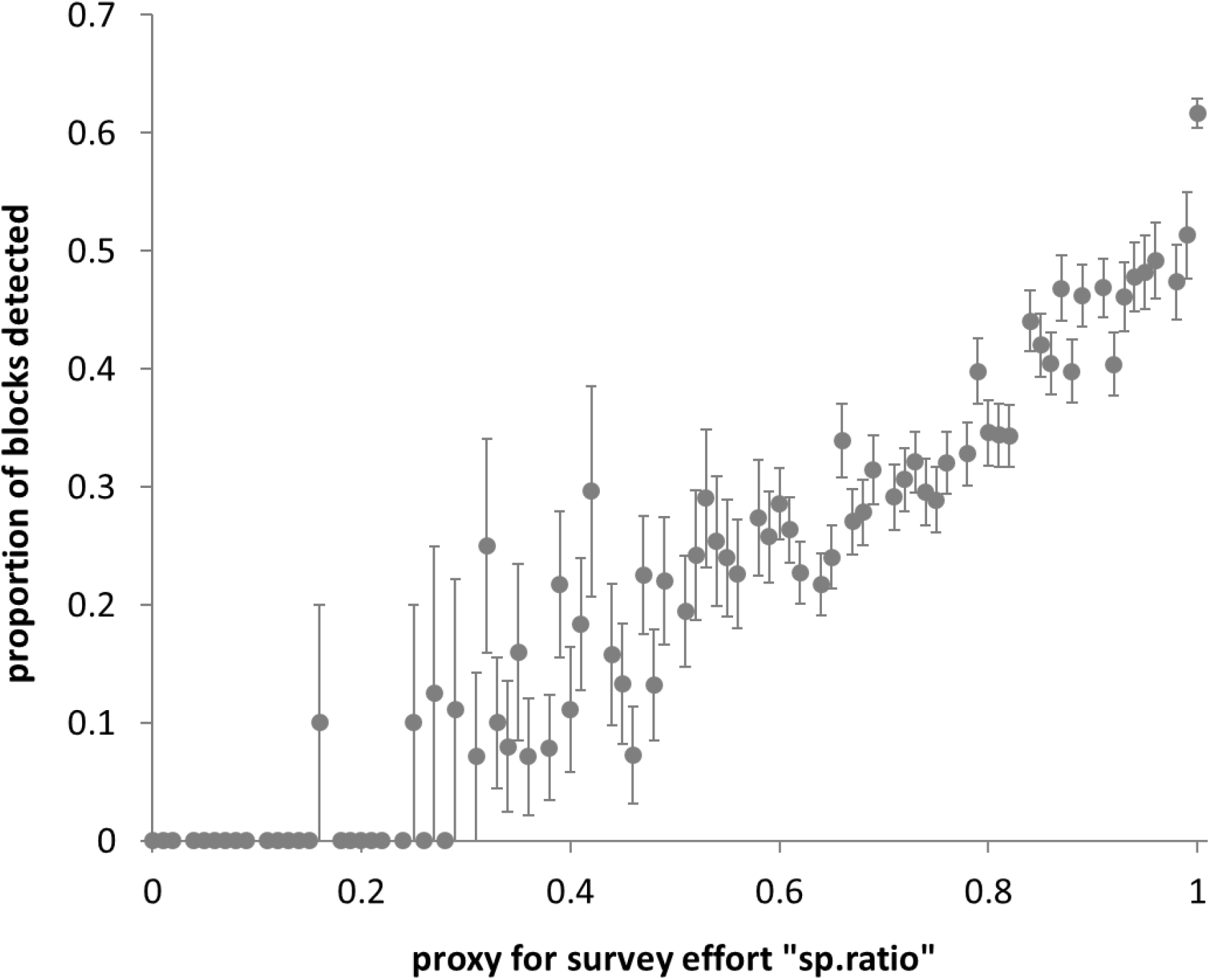
The probability of Broad-winged Hawk detection in an atlas block is approximately linearly correlated with our proxy measure of survey effort: “sp.ratio”, which is the ratio of detected species richness to a hypothetical figure of 75 species. Chart shows relationship for 2nd Atlas data.

We predicted the probability of Broad-winged Hawk presence in each atlas block, for each atlas period, using Bayesian latent Gaussian models fitted using the INLA (Integrated Nested Laplace Approximation) package in Program R (Blangiardo et al. 2013). The probability of Broad-winged Hawk presence in block *O*_*i*_ follows a Bernoulli distribution, and our proxy of survey (sp.ratio) effort is included as a covariate s_*i*_:

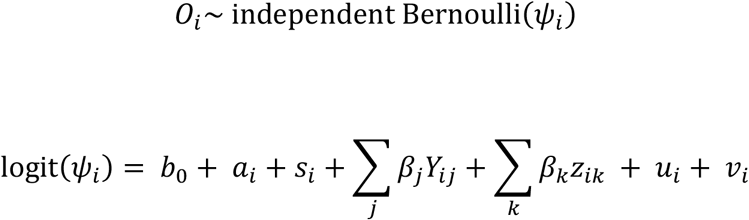

Where *b*_0_ is the intercept, *a* is block area, *Y* are _*j*_ landscape covariates (elevation and latitude) with linear effects *ß*_*j*_, *Z* are _*k*_ land cover covariates (land cover and fragmentation metrics) with linear effects *ß*_*k*_, *u*_*i*_is the spatially structured residual using a Besag-York-Mollie (BYM) specification (Blangiardo and Cameletti 2015), and *v*_*i*_ are unstructured residuals. To account for gaps between blocks within the study area (i.e. non-contiguity of digitized block boundaries along some state lines), we used Voronoi tessellation of atlas blocks as the spatial structure. Block area is included because blocks are not equal size—most blocks along state boundaries are partial blocks.

To determine whether atlas block-level Broad-winged Hawk presence is driven primarily by landscape or land cover, or a combination of the two, we fitted five models:

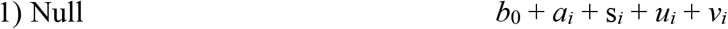

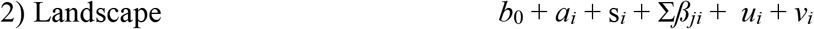

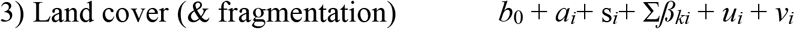

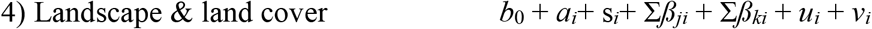

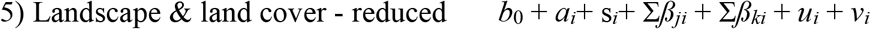

The reduced model (5) was derived heuristically by including all covariates in a non-spatial GLM and selecting only those covariates retained by a forward selection algorithm of the stepAIC function R (Venables and Ripley 2002). Candidate models were assessed using Watanabe-Akaike information criterion (WAIC; (Link and Sauer 2016).

We calculated the difference between the probability of Broad-winged Hawk block occupancy in the 1^st^Atlas and 2^nd^ Atlas, and changes in block-level land cover metrics between the two time periods. We then used the same spatial regression modeling procedure as used to determine the probability of Broad-winged Hawk presence in each atlas period to identify landscape and land cover metrics that were associated with changes in the probability of Broad-winged Hawk presence between atlas periods. Predicted changes in Broad-winged Hawk presence (*δ*_*i*_) were approximately normally distributed (mean -0.128, standard deviation 0.173), so were modeled using Gaussian likelihood model:

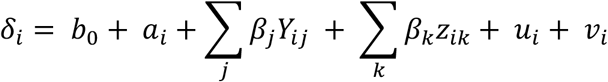

Hence, the model structure is the same as used in the models to predict Broad-winged Hawk distribution for each atlas period.

Because we were interested to see whether changes in Broad-winged Hawk distribution could be separately attributed to land cover change and fragmentation, we added a sixth model that included land cover covariates, but not fragmentation:

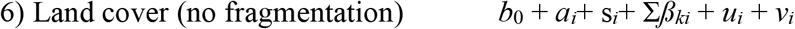

Post-hoc analysis to show the relationships between changes in landscapes (at the block scale) and the predicted change in the probability of Broad-winged Hawk presence (*δ*_*i*_) was done using a GAM smooth from the R package ggplot (Wickham 2011).

### Evaluating Model Performance

We calculated Area under the Receiver Operator Curve (AUC) values to determine the success of predicting Broad-winged Hawk block detections in each atlas period. Because the original data included a large number of false negatives (blocks with little or no survey effort), we calculated AUC for a test sample of 10% of the blocks with comprehensive coverage (75 or more species recorded). Our test sample was 454 for the 1^st^ Atlas and 659 for the 2^nd^Atlas.

## RESULTS

### Changes in Land Cover between Atlas Periods

Changes in land cover reflect a general conversion of forest to other land cover types within the study area. Forested land decreased by 3,584 km^2^ or an average of 0.19 km^2^ per block between the two atlas periods. The number of core patches, area of farmland and area of wetlands also showed an overall decrease between atlas periods. In contrast, the area of urban land increased by 5,657 km^2^, or an average of 0.30 km^2^ per block. The area of barren land and area of water also showed an overall increase between the two atlas periods. There was a net increase in overall land cover diversity.

### Broad-winged Hawk Presence Predictions

In both the 1^st^ Atlas and 2^nd^ Atlas, the best models for predicting the presence of Broad-winged Hawks (based on WAIC) included landscape, land cover, and fragmentation covariates, i.e. model 5 (Table 2). Maps of predicted probability of Broad-winged Hawk presence indicate that the species was significantly under-recorded in both atlas projects in West Virginia, relative to the other states (Figures 2 and 3). AUC values of 0.853 for the 1^st^ Atlas and 0.885 for the 2^nd^ Atlas, suggest that the models were very good at predicting Broad-winged Hawk distributions.

**Table 2.** Model selection for models to predict the species distribution in both atlases, and change between atlases, assessed by change in WAIC relative to a null model. Best models are in bold.

| Model n | $\Delta$ WAIC | | |
| --- | --- | --- | --- |
|  | 1st Atlas<br>(1980s) | 2nd Atlas<br>(2000s) | Change |
| 1 Null | - | - | - |
| 2 Landscape | 779.2 | 737.9 | 5,624.6 |
| 3 Land cover + frag | -980.0 | -1095.7 | **-2,738.3** |
| 4 Landscape + land cover + frag | 244.7 | -145.7 | -961.27 |
| 5 Landscape + land cover + frag (reduced) | **-1010.8** | **-1218.6** | -961.52 |
| 6 Land cover | n.a. | n.a. | -1,553.6 |

**Figure 2.**
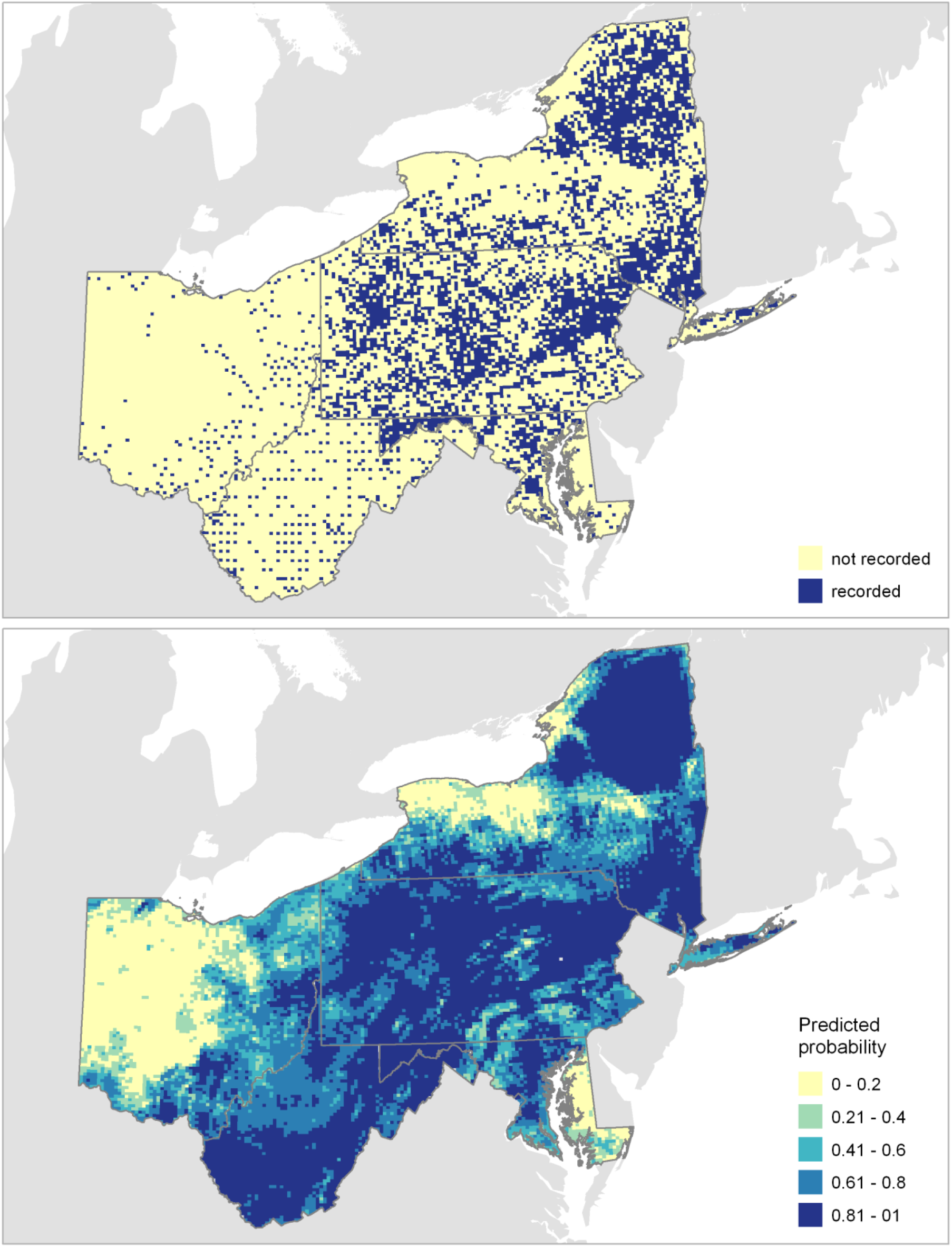
Recorded (top) and predicted (bottom) Broad-winged Hawk distribution in the 1st Atlas period (1980s).

**Figure 3.**
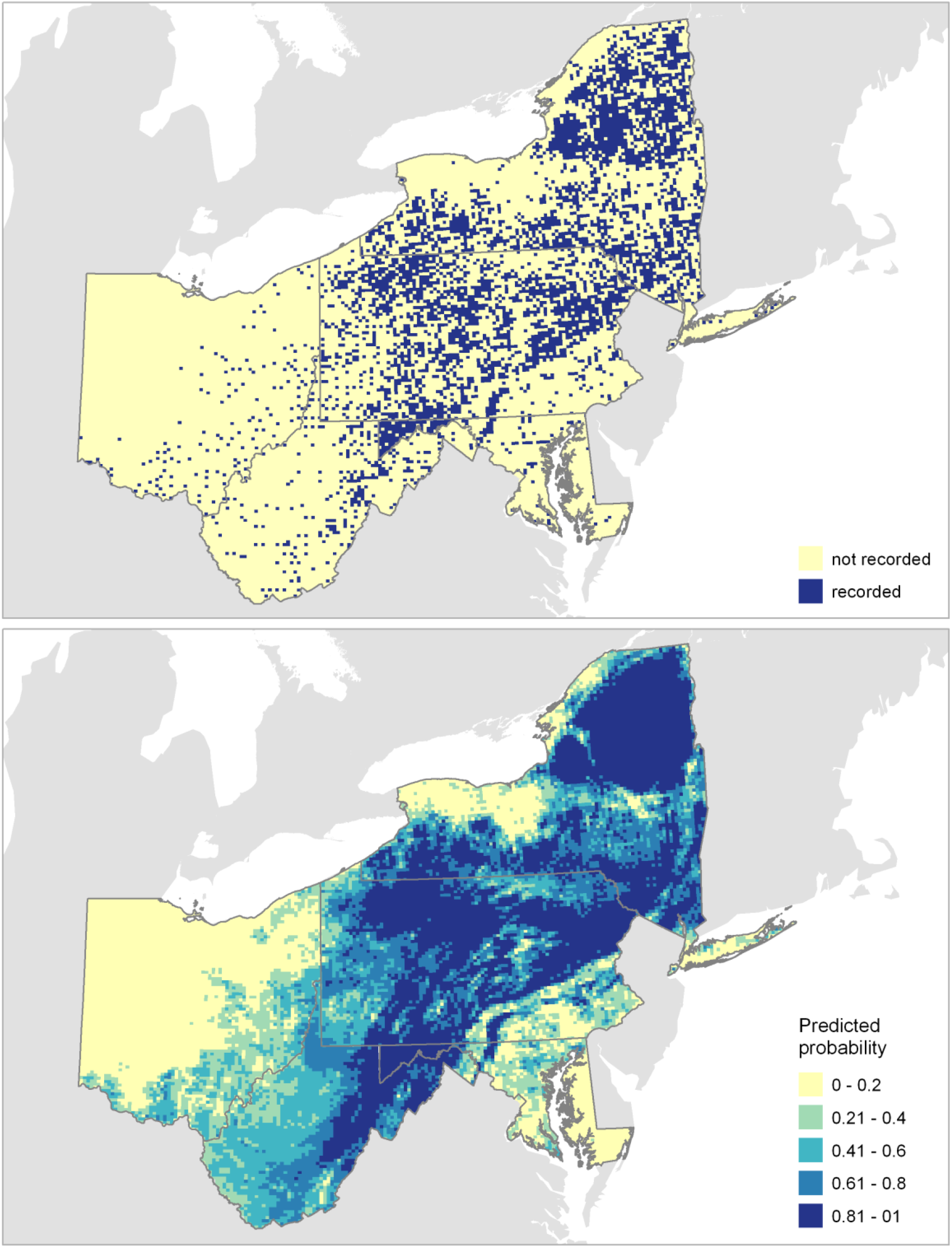
Recorded (top) and predicted (bottom) Broad-winged Hawk distribution in the 2nd Atlas period (2000s).

### Change in Broad-winged Hawk Presence

The best model for predicting changes in Broad-winged Hawk presence including changes in land cover and fragmentation metrics (model 3) but did not include landscape metrics (Table 2). There were positive relationships between predicted changes in Broad-winged Hawk presence and changes in forest area and wetland area, and negative relationships for water, urban, barren and farmland (Table 3). This suggests that Broad-winged Hawks were most likely to have expanded their range into blocks with increasing natural habitat, and more likely to have been lost from blocks with increasing anthropocentric habitat. Increases in the number of core forest patches, largest core patch, core to edge ratio, and habitat diversity were all associated with positive changes in predicted probability of Broad-winged Hawk presence (Table 3). Predicted losses of Broad-winged Hawks were more likely in blocks where core forest declined, the number of core forest patches increased, and the size of the largest forest patch decreased (Figure 5). Losses were also more likely at elevations of less than 500 m (Figure 5).

**Table 3.** Parameter estimates (β) for model of change in Broad-winged hawk distribution between 1^st^ and 2^nd^ Atlases, with standard deviation (SD) and 95% credible intervals (LCI, UCI).

| | $\beta$ | SD | LCI | UCI |
| --- | --- | --- | --- | --- |
| Intercept | -0.2541 | 0.0036 | -0.2612 | -0.2471 |
| block area | 0.0025 | 0.0002 | 0.0022 | 0.0028 |
| $\Delta$ count core forest patches | 0.0005 | 0 | 0.0004 | 0.0005 |
| $\Delta$ ratio of core to edge forest | 0.0002 | 0.0016 | -0.003 | 0.0033 |
| $\Delta$ log(largest forest patch size) | 0.0114 | 0.0008 | 0.0098 | 0.013 |
| $\Delta$ forest area | 0.013 | 0.0009 | 0.0112 | 0.0148 |
| $\Delta$ water area | -0.0074 | 0.005 | -0.0172 | 0.0023 |
| $\Delta$ urban area | -0.0205 | 0.0012 | -0.0229 | -0.0181 |
| $\Delta$ barren area | -0.0087 | 0.0024 | -0.0135 | -0.0039 |
| $\Delta$ farmland area | -0.0198 | 0.001 | -0.0218 | -0.0179 |
| $\Delta$ wetland area | 0.0019 | 0.001 | -0.0001 | 0.0039 |
| $\Delta$ habitat diversity (H) | 0.0284 | 0.0154 | -0.0019 | 0.0587 |

Predicted changes in Broad-winged Hawk presence indicate large-scale losses of this species from northwestern Ohio, the Piedmont region of Maryland and Pennsylvania, and Long Island (NY), and a thinning of the distribution throughout the western half on the northern Appalachian Mountains (Figure 4). Recorded overall change in presence per block showed a reduction of 10.7% across all five states (Table 3). However, our models, which account for changes in survey effort between atlas periods, predict that a range loss (measured in atlas blocks occupied) of 20.7% occurred; ranging from -8.8% in New York to -32.2% in Maryland/DC (Table 4).

**Figure 4.**
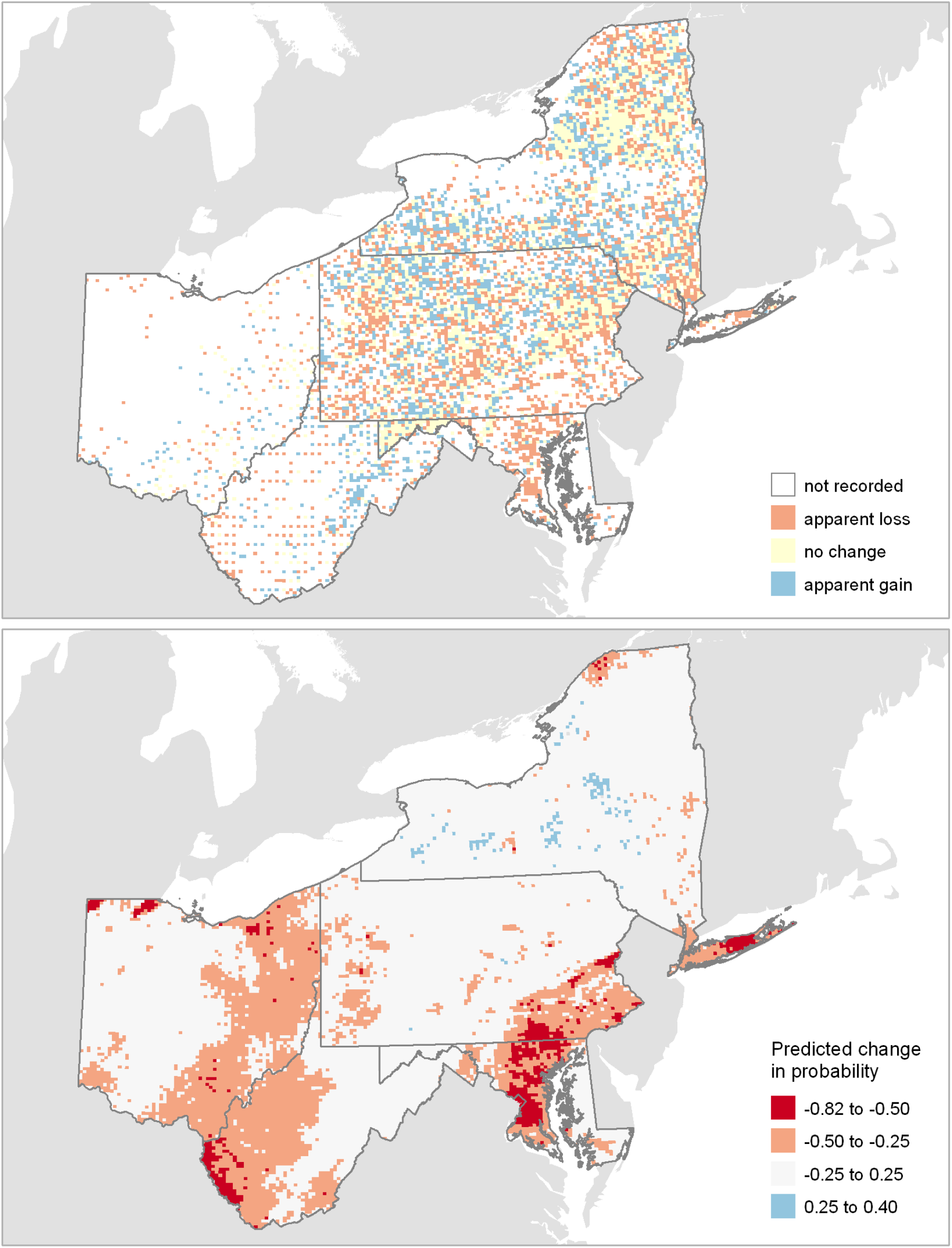
Recorded (top) and predicted (bottom) change in Broad-winged Hawk distribution between the 1^st^ (1980s) and 2^nd^ (2000s) atlas periods. Color symbols: ColorBrewer2.org.

**Figure 5.**
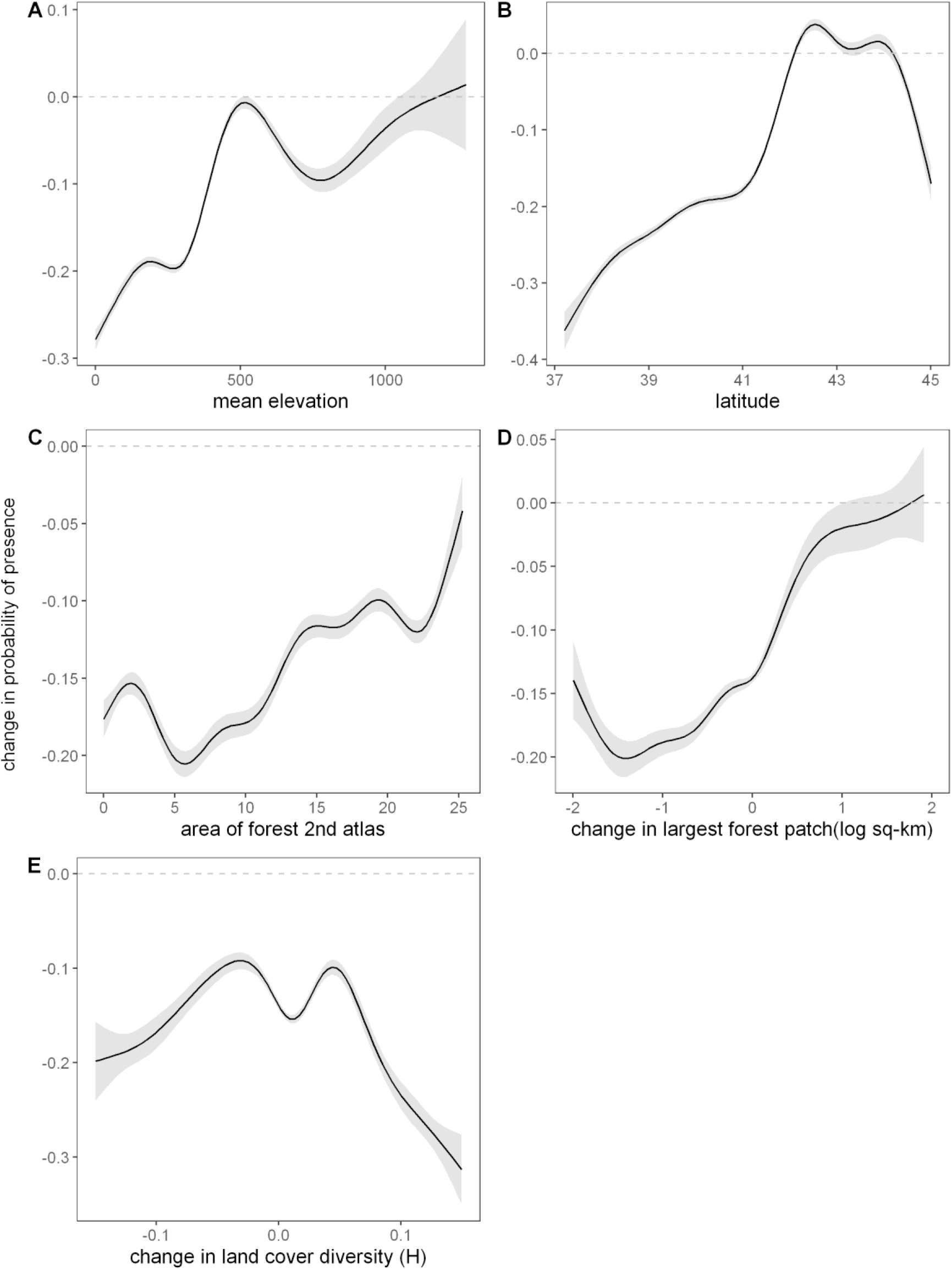
Relationships between landscape characteristics and predicted change in Broad-winged Hawk probability of presence between 1^st^ and 2^nd^ atlas periods. The x-axes of plots D and E exclude extreme outliers (>3 standard deviations from the mean), for clarity.

**Table 4.** Recorded and predicted percentage changes in blocks with Broad-winged Hawks, by state, between 1st Atlases (1980s) and 2nd Atlases (2000s).

|  | Recorded<br>change | Predicted<br>Change |
| --- | --- | --- |
| Maryland/DC | -45.6 | -32.2 |
| New York | +0.9 | -8.8 |
| Ohio | -18.9 | -29.6 |
| Pennsylvania | -16.4 | -20.4 |
| West Virginia | -1.6 | -24.9 |
| All five states | -10.7 | -20.7 |

## DISCUSSION

Our results suggest that the substantial losses of Broad-winged Hawks from areas peripheral to the central Appalachian Mountains since the 1980s is due to land use change. Specifically, we conclude that the increase in anthropocentric non-forest habitats within rapidly developing urban and suburban areas has resulted in the loss of this species from low-elevation areas peripheral to the Appalachian Mountains. Further, the positive relationships between changes in Broad-winged Hawk distribution and indicators of forest contiguity (core forest, core to edge ratio, forest patch size) indicate that forest fragmentation, and not just forest loss, have contributed to the decline in Broad-winged Hawk distribution.

The Broad-winged Hawk may be able to cope with increasing edge forest as long as there are also large tracts of core forest in its range. Rusch and Doerr (1972) report observations of Broad-winged Hawks along edges of large contiguous forest patches in New York. Because Broad-winged Hawks often forage along forest edges, it is possible that they may be finding more food resources with increasing edge forest area. Broad-winged Hawks consume a variety of prey including small mammals as well as reptiles and small birds whose numbers may be increased along forest edges (Rusch and Doerr 1972, McCabe et al. 2019). Increasing edge forest area may not, by itself, be an indication of increasing forest fragmentation, but could also result from reforestation of small patches and woodlots, which would nearly all be classified as edge forest. Similarly, the positive relationship between number of core forest patches and presence of Broad-winged Hawks is open to multiple interpretations. For instance, an increase in number of core patches may be due to increased fragmentation, but could also be due to merging of small forest patches. The positive association of these forest landscape metrics suggests that forest area itself, regardless of whether it is fragmented, is likely the major driver of Broad-winged Hawk distribution at the atlas block scale.

Although this analysis does not definitively determine the environmental drivers related to change in Broad-winged Hawk distribution in specific regions (e.g., the drivers acting in Maryland may be different than drivers of the change observed in southwestern West Virginia), some trends are apparent. Potential impacts of increased urbanization are most apparent in the Piedmont of central and eastern Maryland and southeastern Pennsylvania, as well as northwestern Ohio and Long Island, NY (Figure 4). The Piedmont of Maryland and Pennsylvania is within the Chesapeake Bay watershed, which saw an increase in urban land cover of than 20% between 1990 and 2000, particularly around Washington, DC, Baltimore, MD and Philadelphia, PA (Jantz et al. 2005). Broad-winged Hawks appear to be more sensitive to urbanization than other forest raptors, such as Red-shouldered and Cooper’s hawks (Dykstra et al. 2000, Ellison 2010, Rodewald et al. 2016). Nesting Broad-winged Hawks were reported to avoid human dwellings in Ontario (Armstrong and Euler 1983) and to occur more often in undisturbed forests (Devaul 1990), which supports our findings. Forest fragmentation is associated with higher rates of nest predation for songbirds (Thompson III et al. 2002). Higher nest predation rates also could explain why Broad-winged Hawks are less abundant in fragmented landscapes. Thus, fragmentation and more permanent replacement of forest by landcover types such as suburban development may have negative consequences to Broad-winged Hawks. Indeed, the majority of landscape metrics associated with loss of Broad-winged Hawks were related to changes in non-forest land cover types. These associations suggest that increases in non-forest land cover in the study region are contributing to range losses of the Broad-winged Hawk.

It is likely that the shift in range is not attributed to solely one agent; rather a combination of habitat loss, fragmentation, prey availability, predator-prey dynamics, climate, or other biotic factors that may have an influence on habitat suitability. It is important to note that the relative contribution of each of the landscape metrics was not differentiated in this analysis and further analysis of the degree of influence may be beneficial for better understanding the principal drivers of distribution change.

### Landscape Characteristics

The results of our study suggest that changes in Broad-winged Hawk distribution are not directly related to elevation and latitude. Given that the areas from which this species has retreated are predominantly at low elevations, and the fact that the species has shown less net range change in the north (specifically New York), this result is somewhat surprising. However, it is possible that our models failed to tease out the separate effects of topography and land cover due to the intercorrelation between them. Hence, we do not consider our study to be sufficient to rule out climate change as a contributory driver of the range loses shown.

### Limitations

The mismatch of time periods between NLCD data and the 1st Atlas data (1992 land cover versus 1980s bird data) introduces some bias to the analysis, namely that change in land-cover may have been underestimated. The secretive nature of the Broad-winged Hawk may have led to a large number of false negatives in the atlas data set. Although we attempted to overcome this issue assuming a linear relationship between effort (number species reported in each block) and the probability of detecting Broad-winged Hawks, we acknowledge that this approach is somewhat simplistic, and that the actual relationship between species richness and hawk detection may be non-linear, and may vary regionally. While these limitations are not trivial, the tremendous amount of data available on this species through the combined ten atlas projects confers some confidence that the spatial patterns observed are genuine.

Many questions remain regarding the area of land that is necessary for persistence of Broad-winged Hawks, the most relevant drivers of the range shift, and the impacts that shifts may have on Broad-winged Hawk populations. Further analysis is warranted to support conclusions of the drivers defined in this study. Additionally, similar methodology could be applied to other forest-interior species, particularly those in decline, to determine the key landscape metrics related to their breeding range extent and may be especially important for species that may be in decline.

### Conservation Implications and Recommendations

Preserving large tracts of contiguous forest will become increasingly important to support forest dependent species. A shift in range and decrease of suitable habitat may lead to increased competition between other larger hawk species and the Broad-winged Hawk for nesting sites (Ellison 2010). Since there is an association with large areas of core forest, these areas should continue to be the focus of conservation strategies. Reducing the conversion of forest to non-forested land is also an important and influential component of habitat conservation for this and many other forest species. A better understanding of how changes in the landscape are altering the habitat choices of the Broad-winged Hawk, as well as other species, is important for ensuring that these habitats remain present in future landscapes. Determination of regions where the greatest threats occur may help resource managers to better define important bird conservation priorities by region, and identify areas to preserve. Increasing connectivity of forest in urbanizing regions of the east, through forest and riparian corridor restoration as well as maintaining-forested neighborhoods, may improve available habitat and ameliorate the patterns observed in this study.

## ACKNOWLEDGEMENTS

This study was only possible due to the efforts of many thousands of citizen scientists who contributed to the 10 bird atlas projects over three decades. We thank Richard Bailey (West Virginia DNR), Walter Ellison (Washington College), John Ozard (NYSDEC Bureau of Wildlife) for sharing Broad-winged Hawk occupancy and species data for the respective state atlas projects. This analysis was undertaken for an undergraduate honors thesis in the Environmental Studies Department at Gettysburg College, who we thank for financial and IT support. This is Hawk Mountain conservation science contribution number (*number to be provided for an accepted MS*).

## Appendix

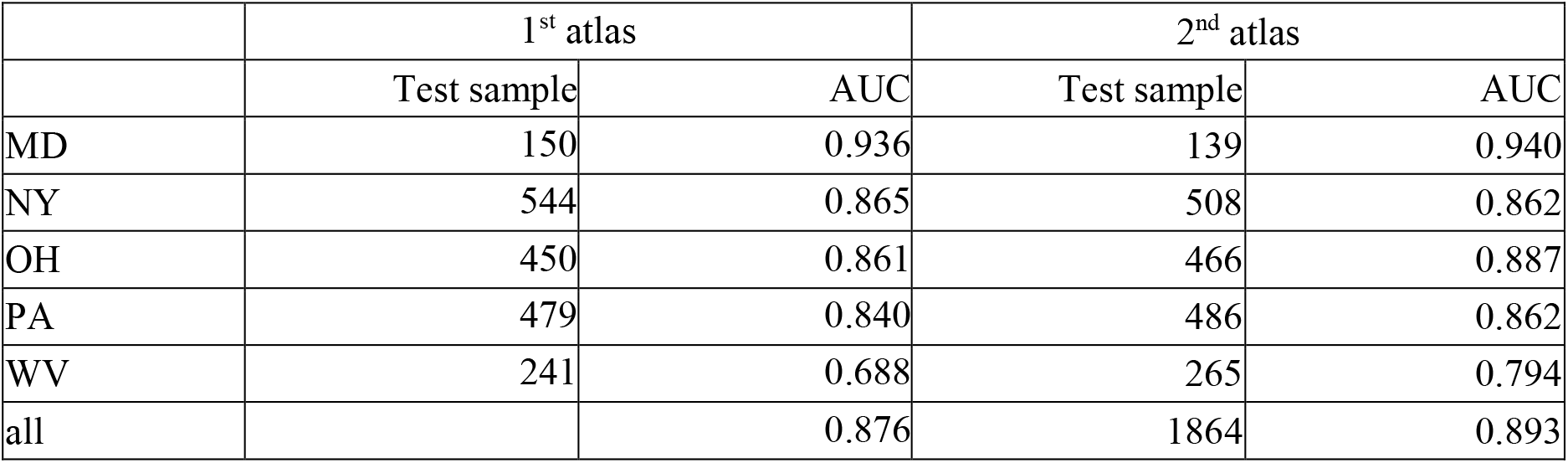

